# Persistence of SARS-CoV-2 specific B- and T-cell responses in convalescent COVID-19 patients 6-8 months after the infection

**DOI:** 10.1101/2020.11.06.371617

**Authors:** Natalia Sherina, Antonio Piralla, Likun Du, Hui Wan, Makiko Kumagai-Braesh, Juni Andréll, Sten Braesch-Andersen, Irene Cassaniti, Elena Percivalle, Antonella Sarasini, Federica Bergami, Raffaella Di Martino, Marta Colaneri, Marco Vecchia, Margherita Sambo, Valentina Zuccaro, Raffaele Bruno, Tiberio Oggionni, Federica Meloni, Hassan Abolhassani, Federico Bertoglio, Maren Schubert, Miranda Byrne-Steele, Jian Han, Michael Hust, Yintong Xue, Lennart Hammarström, Fausto Baldanti, Harold Marcotte, Qiang Pan-Hammarström

## Abstract

**Background:** The longevity of the immune response against SARS-CoV-2 is currently debated. We thus profiled the serum anti-SARS-CoV-2 antibody levels and virus specific memory B- and T-cell responses over time in convalescent COVID-19 patients.

**Methods:** A cohort of COVID-19 patients from the Lombardy region in Italy who experienced mild to critical disease and Swedish volunteers with mild symptoms, were tested for the presence of elevated anti-spike and anti-receptor binding domain antibody levels over a period of eight months. In addition, specific memory B- and T-cell responses were tested in selected patient samples.

**Results:** Anti-SARS-CoV-2 antibodies were present in 85% samples collected within 4 weeks after onset of symptoms in COVID-19 patients. Levels of specific IgM or IgA antibodies declined after 1 month while levels of specific IgG antibodies remained stable up to 6 months after diagnosis. Anti-SARS-CoV-2 IgG antibodies were still present, though at a significantly lower level, in 80% samples collected at 6-8 months after symptom onset. SARS-CoV-2-specific memory B- and T-cell responses were developed in vast majority of the patients tested, regardless of disease severity, and remained detectable up to 6-8 months after infection.

**Conclusions:** Although the serum levels of anti-SARS-CoV-2 IgG antibodies started to decline, virus-specific T and/or memory B cell responses increased with time and maintained during the study period (6-8 months after infection).

**Funding:** European Union’s Horizon 2020 research and innovation programme (ATAC), the Italian Ministry of Health, CIMED, the Swedish Research Council and the China Scholarship Council.

## Introduction

The emergence and spread of a novel coronavirus, SARS-CoV-2, has led to a pandemic with a major impact on global health. Currently, the immunological features associated with severity of the disease or protection remains largely unknown and whether the antibody titer is a marker for protective immunity against SARS-CoV-2 is currently debated. A robust adaptive immune response with presence of spike-specific neutralizing antibodies, memory B cells and circulating follicular helper T cells have been found in patients who have recovered from infection ^1 2 3^. It is however still unclear how long the adaptive immunity to SARS-CoV-2 lasts after the natural infection. A relationship between a humoral immune response to SARS-CoV-2 infection and protection against reinfection has been shown in rhesus macaques ^4^ but remains to be determined in humans. While a recent study in Iceland showed that the antibody response was maintained in 90% of convalescent patients for more than four months after onset of disease ^5^, other studies suggest a rapid decay of anti-SARS-CoV-2 IgG in individuals with mild illness ^6 7^. Nevertheless, long-lived memory T and B cells could potentially be present and reactivated following a second exposure, thus providing immune protection.

The SARS-CoV-2 virus genome encodes four major structural proteins including the spike protein (S), nucleoprotein (N), membrane protein (M), envelop (E) and other membrane proteins (ORF3a, ORF7a) ^8^. The primary target for an antibody-mediated response on the surface of SARS-CoV-2 virions is the homotrimeric S protein ^9^. Antibodies targeting the S protein and its receptor-binding domain (RBD) are therefore of particular interest to combat the infection and a large range of RBD-specific monoclonal antibodies isolated from convalescent patients neutralize the virus both *in vitro* and in animal models ^2 10 11^.

Studies on the longevity of the adaptive immune response in convalescent COVID-19 patients may facilitate understanding of how immune protection develops and persists during the natural course of SARS-CoV-2 infection and provide useful information for the development of vaccines against this newly emerging virus. In this study, we aimed to assess the dynamics and longevity of the SARS-CoV-2-specific immune responses in COVID-19 patients with broad spectrum of disease scores. Levels and Ig class of SARS-CoV-2-specific antibodies and development of memory B and T cells were evaluated in samples collected from 88 patients at different time points during a period of 8 months following initial symptoms.

## Methods

### Patients and sample collection

Screening of COVID-19 patient donors and sample collection were conducted at the Fondazione IRCCS Policlinico San Matteo in Pavia, Italy, a designated medical institution for COVID-19. Study inclusion criteria included subjects over 18 years of age, who were willing and able to provide informed consent, confirmed positivity of SARS-CoV-2 by real-time RT-PCR targeting the *E* and *RdRp* genes according to Corman *et al*. protocols ^12^ and monitored until two subsequent samples with negative results. Between February 28 and October 10, 2020, 78 COVID-19 patients were recruited. Forty-seven donors had blood drawn at one single time point ranging from 7 to 240 days after symptom onset while 28 and 3 donors had blood taken at two or three time points, respectively. Disease severity was defined as mild (non-hospitalized), moderate (hospitalized, with lower respiratory tract infection, with dyspnea or not, but without oxygen support), severe (infectious disease/sub intensive ward with a need for oxygen and/or positive chest computed tomography scan, severe lower tract infections, with any oxygen support) and critical (intensive care unit (ICU) patients, intubated or with extracorporeal membrane oxygenation procedures).

The demographic and clinical characteristics of the patients are summarized in Table 1 and detailed in S1. The patients had a median age of 63 years (range 32-89) with 45 (58%) males and 33 (42%) females. The degree of clinical severity of COVID-19 in cohort was mild (n=5), moderate (n=16), severe (n=52) and critical (n=5). The most common underlying diseases were hypertension 40/78, 51%), diabetes (16/78, 21%), heart disease (12/78, 15%) and obesity (10/78, 13%). The study was performed under the approval of the Institutional Review Board of Policlinico San Matteo (protocol number P_20200029440).

**Table 1.** Summary of demographic and clinical characteristics in COVID-19 positive individuals

|  | Italian cohort |  | Swedish cohort |  |
| --- | --- | --- | --- | --- |
| Demographic |  |  |  |  |
| Number | 78 |  | 10 |  |
| Age, median (range) | 63.0 (32-89) |  | 53.5 (29-75) |  |
| Male | 58% | (45/78) | 50% | (5/10) |
| Female | 42% | (33/78) | 50% | (5/10) |
| Disease severity |  |  |  |  |
| Mild | 6% | (5/78) | 100% | (10/10) |
| Moderate | 21% | (16/78) | 0% | (0/10) |
| Severe | 67% | (52/78) | 0% | (0/10) |
| Critical | 6% | (5/78) | 0% | (0/10) |
| Symptoms |  |  |  |  |
| Fever | 96% | (75/78) | 40% | (4/10) |
| Cough | 67% | (52/78) | 40% | (4/10) |
| Dyspnea | 47% | (37/78) | 0% | (0/10) |
| Asthenia | 12% | (9/78) | 60% | (6/10) |
| Diarrhea | 9% | (7/78) | 0% | (0/10) |
| Anosmia | 4% | (3/78) | 50% | (5/10) |
| Hypoxia | 1% | (1/78) | 0% | (0/10) |
| Myalgia | 1% | (1/78) | 50% | (5/10) |
| Past medical history |  |  |  |  |
| Hypertension | 51% | (40/78) | 0% | (0/10) |
| Diabetes | 21% | (16/78) | 0% | (0/10) |
| Heart diseases | 15% | (12/78) | 0% | (0/10) |
| Obesity | 13% | (10/78) | 0% | (0/10) |
| HCV | 10% | (8/78) | 0% | (0/10) |
| Lung diseases | 5% | (4/78) | 0% | (0/10) |
| Tumor | 5% | (4/78) | 0% | (0/10) |
| Other comorbidities | 41% | (32/78) | 10% | (1/10) |
| More than one comorbidity | 49% | (38/78) | 0% | (0/10) |
| More than two comorbidities | 29% | (23/78) | 0% | (0/10) |
| Severity |  |  |  |  |
| Oxygen therapy | 73% | (57/78) | 0% | (0/10) |
| Intensive care unit (ICU) | 6% | (5/78) | 0% | (0/10) |

Twelve samples from 10 volunteers from Sweden (median age of 54 years, range 29-75) who had tested PCR- or serology-positive for SARS-CoV-2 and experienced mild symptoms were also included (Table 1 and S1). Blood samples were collected at 60-238 days after onset of symptoms. The study was approved by the ethics committee in Stockholm.

In addition, serum samples from 108 individuals (16 to 80 years of age), collected before the SARS-CoV-2 pandemic (1995 to 2005) were used as historical negative controls for the ELISA and PBMCs from four healthy controls (median age 41 years, range 39-50) and seven additional buffy coats collected in Sweden before the SARS-CoV-2 pandemic (2011-January 2020) were included as negative controls for the B- and T-cell assays. Patients and samples tested in different assays were summarized in a flow chart (Figure 1).

**Figure 1.**
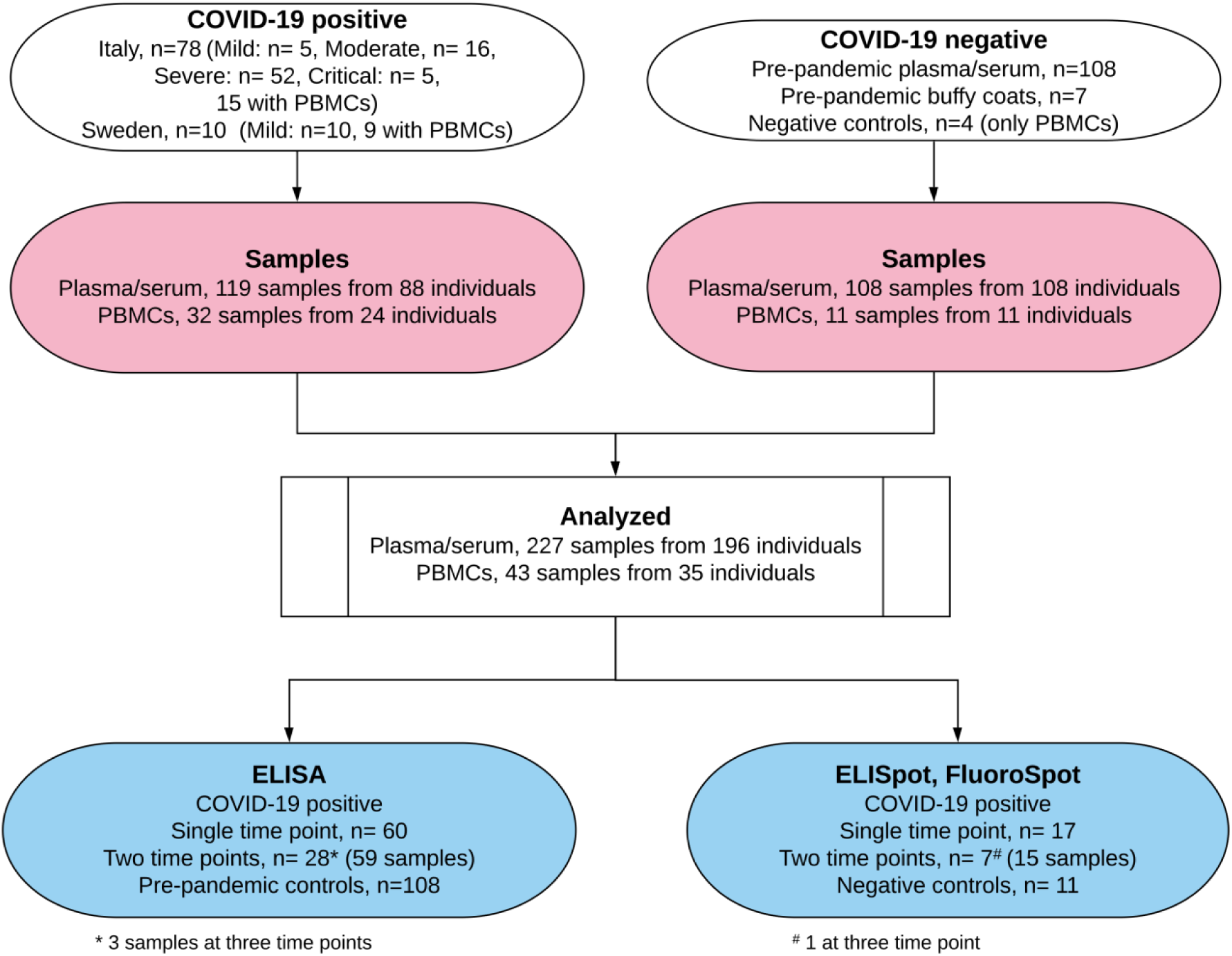
Flow chart illustrating the study design.

### Detection of antibodies specific to SARS-CoV-2

RBD-His protein was expressed in Expi293 cells and purified on Ni-NTA resin (#88221, Thermofisher) followed by size-exclusion chromatography on a Superdex 200 gel filtration column in PBS ^13^. S1-S2-His (referred as S) protein was expressed baculovirus-free in High Five insect cells ^14^ and purified on HisTrap excel column (Cytiva) followed by preparative size exclusion chromatography on 16/600 Superdex 200 kDa pg column (Cytiva) ^15^.

High-binding Corning Half area plates (Corning #3690) were coated over night at 4°C with S or RBD protein (1.7 μg/ml for IgG and IgM; 2.0 μg/ml for IgA) in PBS; washed three times in PBS-Tween (0.05%) and blocked with 2% BSA in PBS for 1h at room temperature. Serum or plasma diluted 1:6400 (S IgG), 1:3200 (S IgM), 1:1600 (S IgA; RBD IgG, IgA, IgM) in 0.1% BSA in PBS, was incubated for 1.5h at room temperature. Plates were then washed and incubated for 1h at room temperature with horseradish peroxidase (HRP)-conjugated goat anti-human IgG (Invitrogen #A18805), goat anti-human IgM (Invitrogen #A18835), or goat anti-human IgA (Jackson #109-036-011) (all diluted 1:15 000 in 0.1% BSA-PBS). Bound antibodies were detected using tetramethylbenzidine substrate (Sigma #T0440). The color reaction was stopped with 0.5M H_2_SO_4_. Absorbance was measured at 450nm. Antibody levels were presented as arbitrary units (AU/ml), based on a standard curve made from a serially diluted highly positive serum pool. A cut-off value for antibody positivity was defined for each antigen and isotype using receiver operating characteristic (ROC) curves, based on the antibody responses in historical controls (n=108) and COVID-19 patients (55 samples collected within 7-28 days after symptom onset). The cut-off value for positivity was set at >0.03 AU/ml for anti-S IgG, >0.5 AU/ml for anti-S IgA, >2.5 AU/ml for anti-S IgM, >14.8 AU/ml for anti-RBD IgG, >0.08 AU/ml for anti-RBD IgA, and >8.4 AU/ml for anti-RBD IgM, giving a specificity of 97% for IgG, 99% for IgA and 96% IgM. A previously described microneutralization assay ^16 17^ was used to determine the titers of SARS-CoV-2 neutralizing antibodies (NT-abs) in 37 samples. The neutralizing titer was the maximum dilution giving a reduction of 90% of the cytopathic effect.

### Isolation of PBMCs and RNA

PBMCs were isolated from blood or buffy coat samples by standard density gradient centrifugation using Lymphoprep (Axis-Shield) and were cryopreserved and stored in liquid nitrogen until analysis. Total RNA was extracted from PBMCs by using RNeasy mini kit according to the manufacturer’s protocol (Qiagen).

### Enumeration of B cells secreting IgG antibodies specific for SARS-CoV-2 RBD and T cells secreting IFN-γ and IL-2 in response to SARS-CoV-2 peptides

PBMCs were incubated for four days in RPMI-1640 medium with 10% FCS, supplemented with the TLR7 and TLR8 agonist imidazoquinoline resiquimod (R848, 1 µg/ml; Mabtech AB, Nacka, Sweden), and recombinant human IL-2 (10 ng/ml) for stimulation of memory B cells ^18^. The ELISpot plates pre-coated with capturing monoclonal anti-human IgG antibodies were incubated with a total of 300 000 or 30 000 pre-stimulated cells per well for detection of RBD-specific IgG and total IgG secreting cells, respectively. The number of B cells secreting IgG antibodies specific for SARS-CoV-2 RBD and cells secreting IgG (total IgG) were measured using the Human IgG SARS-CoV-2 RBD ELISpot^PLUS^ (ALP) kit according to the manufacturer’s protocol (Mabtech AB).

IFN-γ and IL-2 secreting T cells were detected using Human IFN-γ/IL-2 SARS-CoV-2 FluoroSpot^PLUS^ kits according to the manufacture’s protocol (Mabtech AB). The plates pre-coated with capturing monoclonal anti-IFN-γ and anti-IL-2 were incubated overnight in RPMI-1640 medium containing 10% FCS supplemented with a mixture containing the SARS-CoV-2 peptide pool (scanning or defined pools), anti-CD28 (100 ng/ml) and 300 000 cells per well in a humidified incubators (5% CO_2_, 37°C).

The SARS-CoV-2 S1 scanning pool contains 166 peptides from the human SARS-CoV-2 virus (#3629-1, Mabtech AB). The peptides are 15-mers overlapping with 11 amino acids, covering the S1 domain of the S protein (amino acid 13-685). The SARS-CoV-2 S N M O defined peptide pool contains 47 synthetic peptides binding to human HLA, derived from the S, N, M ORF3a and ORF7a proteins (#3622-1, Mabtech AB) ^19^. The SARS-CoV-2 S2 N defined peptide pool contains 41 synthetic peptides binding to human HLA derived from the S and N proteins of the SARS-CoV-2 virus (#3620-1, Mabtech AB) ^20^. Results of ELISpot and Fluorospot assays were evaluated using an IRIS-reader and analyzed by IRIS software version 1.1.9 (Mabtech AB). The results were expressed as the number of spots per 300 000 seeded cells after subtracting the background spots of the negative control. The cut-off value was set at the highest number of specific B- and T-cell spots from the negative controls.

### Statistical analysis

Mann-Whitney U test was used for comparisons between groups in anti-SARS-CoV-2 antibody levels and numbers of specific memory B- and T-cells. Correlation analysis was performed using Spearman’s rank correlation. A Wilcoxon signed-rank test was used for comparison paired samples. All analyses and data plotting were performed using GraphPad or R version 3.6.1.

## Results

### The dynamics of the anti-SARS-CoV-2 antibody response in COVID-19 patients

In order to evaluate the antibody response to SARS-CoV-2, 119 serum or plasma samples from 88 COVID-19 patients (78 from Italy and 10 from Sweden; Figure 1, Table 1 and Table S1) were tested by an in-house enzyme-linked immunosorbent assay (ELISA) for the presence of anti-S and anti-RBD antibodies. First, we examined the SARS-CoV-2-specific IgG, IgA, and IgM antibodies in 55 samples from COVID-19 patients collected during early phases of recovery (between 7-28 days after onset of disease symptoms) and 108 historical controls (samples collected prior to the SARS-CoV-2 pandemic). Significantly higher levels of anti-S and anti-RBD IgG, IgA and IgM antibodies (p<0.0001 for all groups) were detected in patients as compared to historical controls (Fig. 2A-C). Anti-S IgG, IgA and IgM levels were increased in 85%, 78% and 76% of patients, respectively. A similar proportion of patients had elevated anti-RBD IgG, IgA and IgM levels with 85%, 62% and 67% being positive, respectively. Titers of anti-S and anti-RBD antibodies were highly correlated for all isotypes (r=0.95 for IgG, r=0.77 for IgA, r=0.88 for IgM) (Fig. 2D-F). SARS-CoV-2 NT-ab titers, measured by microneutralization test, correlated with levels of anti-S IgG (r=0.44), anti-S IgA (r=0.34) and anti-S IgM (r=0.53), as well as with levels of anti-RBD IgG (r=0.39) and anti-RBD IgM (r=0.40) (data not shown).

**Figure 2.**
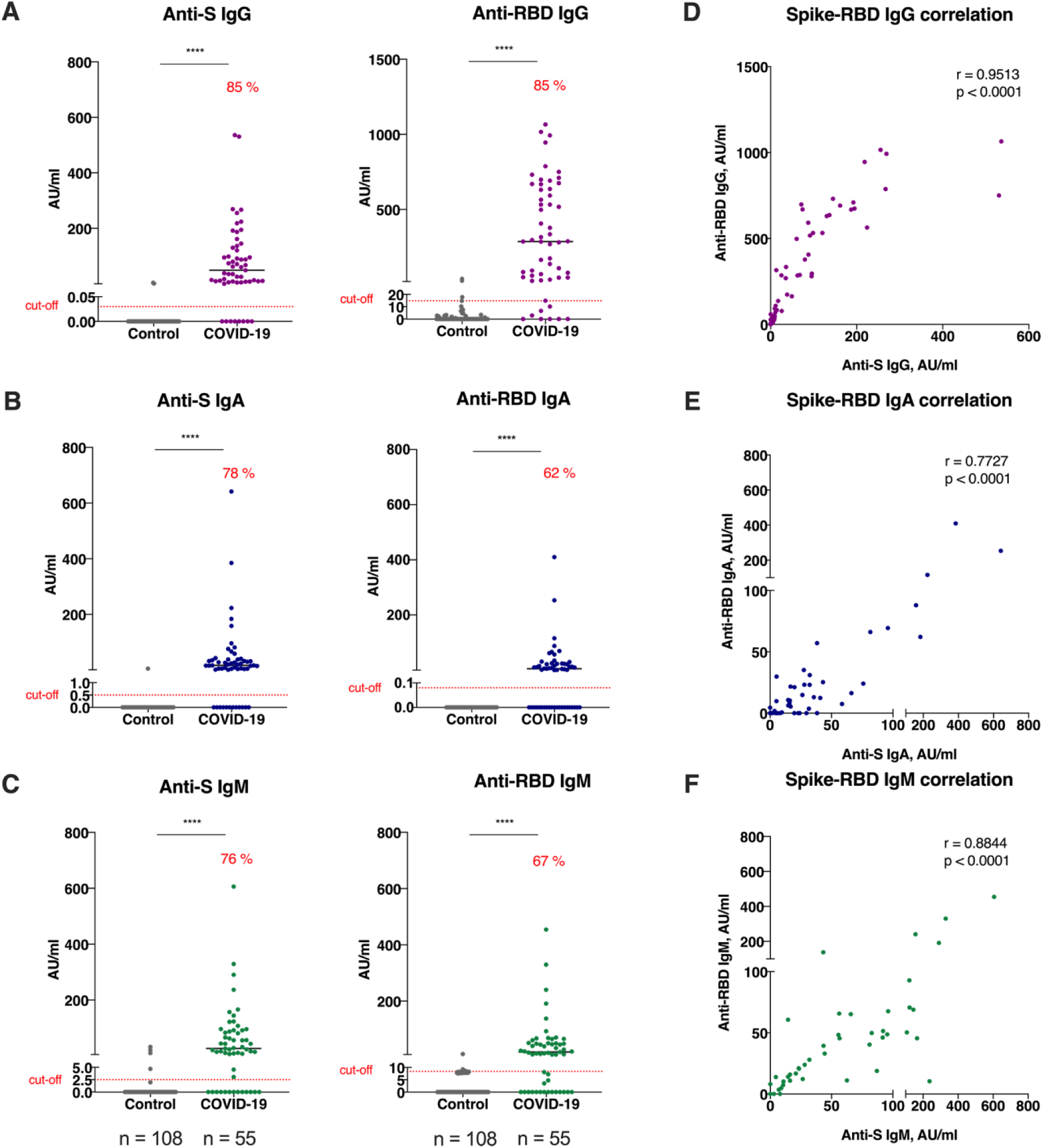
Anti-SARS-CoV-2 antibody response in COVID-19 patients. Levels of anti-S and anti-RBD IgG, IgA and IgM antibodies in historical controls and COVID-19 patients (**A, B, C**). Correlation between titers of anti-S and anti-RBD IgG, IgA, and IgM (**D, E, F**). Symbols represent individual subjects; horizontal black lines indicate the median. The dashed red line indicates the cut-off value for elevated anti-S and anti-RBD antibody levels (0.03 and 14.81 AU/ml for IgG, 0.5 and 0.08 AU/ml for IgA, 2.5 and 8.4 AU/ml for IgM, respectively) defined using receiver operating characteristic (ROC) curves, based on the antibody responses in historical controls (n=108) and COVID-19 patients (n=55). Percentages in graphs A, B and C show the frequency of antibody positive individuals. Statistical significance determined by a non-parametric Mann-Whitney U test (A, B, C), correlation analysis was performed by Spearman’s rank correlation (D, E, F). ****p<0.0001

Based on the symptoms presented at the time of COVID-19 diagnosis, patients were given a disease score ranging from mild, moderate, severe to critical (see MM). Analysis of anti-SARS-CoV-2 antibody levels did not show statistically significant difference in patients presented with severe and critical disease scores compared to mild or moderate disease groups (p=0.1444 and p=0.2943 for anti-S IgG, p=0.1203 and p=0.4672 for anti-RBD IgG, respectively) (Fig. S1A-B). Notably, in six patients (11%), rather low levels or even an absence of anti-S and anti-RBD IgG, IgA and IgM antibodies were observed. Sera from these individuals were obtained at median day 10.5 after symptom onset (range 7-22). A second sample taken at later time point was available for two of these patients (86 and 226 days after symptom onset), where both had become anti-S and anti-RBD IgG positive. For the other four samples (7%), no second sample was available for analysis. These four individuals had a higher median age (79 years) compared to the median age for the entire patient cohort (62 years), had severe disease scores, and two of them later died from COVID-19 complications. When patients were divided based on sex, no statistically significant differences were observed in anti-S and anti-RBD antibody levels for all isotypes, except for anti-RBD IgG, where significantly higher levels were present in males compared to females with severe/critical disease scores (p=0.0306) (Fig. S1C-D).

To examine the longevity of the anti-SARS-CoV-2 antibody response, we subsequently analyzed all 119 samples from 88 patients collected at different time points (7 to 240 days after symptom onset). Anti-S and anti-RBD antibody levels were significantly increased already 7 to 14 days after symptom onset (p<0.0001 for all isotypes) and reached maximum at day 15 to 28 (Fig. 3) (days 7-14 to days 15-28, p=0.0159 and p=0.0049 for IgG, p=0.0080 and p=0.0029 for IgA, p=0.0087 and p=0.0309 for IgM, respectively). After day 28, a significant decrease in anti-S and anti-RBD IgA and IgM antibody levels was observed (days 15-28 to days 29-90, p=0.0184 for anti-RBD IgA; days 15-28 to days 91-180, p<0.0001 for anti-S and anti-RBD IgA and IgM; days 15-28 to days 181-240, p<0.0001 for anti-S and anti-RBD IgA and IgM). No significant decrease in anti-S and anti-RBD IgG levels was present by days 91-180 (days 15-28 to days 91-180, p=0.1847 and p=0.0544, respectively), however a significant decline was observed by days 181-240 (days 15-28 to days 181-240, p=0.0003 and p=0.0002, respectively). Importantly, a prominent anti-SARS-CoV-2 IgG response was still present in 80% (12/15) of the patients who were followed 181-240 days after onset of symptoms. These patients had mild (n=5), moderate (n=2) and severe (n=5) disease scores at time of the diagnosis.

**Figure 3.**
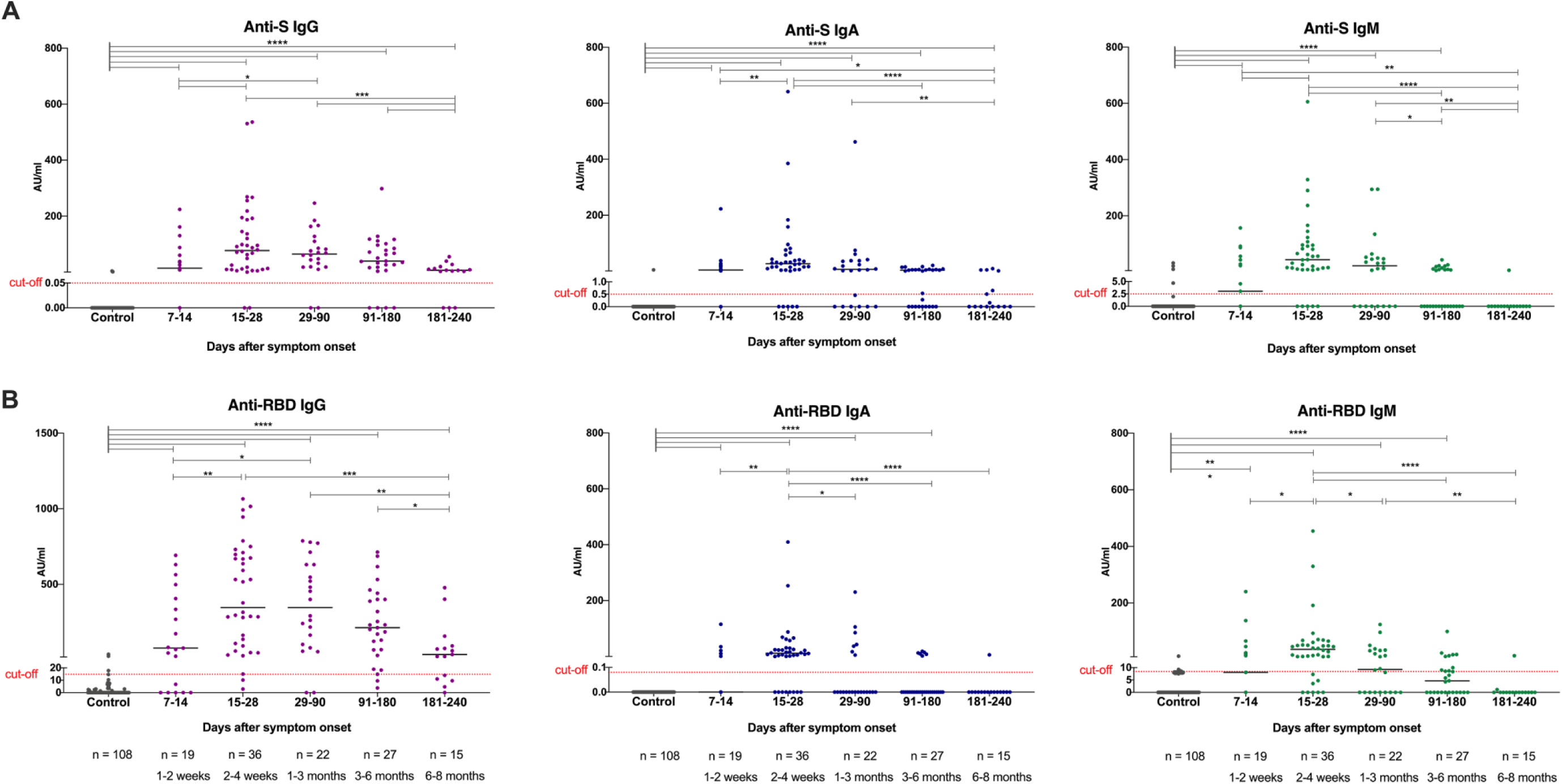
Longevity of the anti-SARS-CoV-2 antibody response in COVID-19 patients. Anti-S and anti-RBD IgG, IgA and IgM antibody response in COVID-19 patients during the time following the diagnosis and recovery (A, B). In total, 119 samples were collected from 88 patients. Samples were taken at 5 study periods: 1-2 weeks (n=19), 2-4 weeks (n=36), 1-3 months (n=22), 3-6 months (n-27), and 6-8 months (n=15) after symptom onset. Symbols represent individual subjects; horizontal black lines indicate the median. The dashed red line indicates the cut-off value for elevated anti-S and anti-RBD antibody levels (0.03 and 14.81 AU/ml for IgG, 0.5 and 0.08 AU/ml for IgA, 2.5 and 8.4 AU/ml for IgM, respectively) defined using receiver operating characteristic (ROC) curves, based on the antibody responses in historical controls (n=108) and COVID-19 patients (n=55). Statistical significance determined by a non-parametric Mann-Whitney U test. *p≤0.05, **p≤0.001, ***p≤0.001, ****p<0.0001

To further evaluate the dynamics of the anti-SARS-CoV-2 antibody response, we compared the antibody levels in paired samples from twenty-seven patients. The first sample was taken at median day 21 (range 7-64) after onset of symptoms and the paired second sample was taken at median day 126 (range 57-234). This analysis showed a significant decrease in anti-S IgA and IgM levels (p=0.0008 and p<0.0001, respectively), and a significant decrease in anti-RBD IgA and IgM levels (p=0.0052 and p=0.0002, respectively) with time (Fig. 4). No significant decline in anti-S and anti-RBD IgG levels was observed (p=0.1551). Taken together, these data suggest that an anti-SARS-CoV-2 antibody response was induced in a majority of COVID-19 patients in our study cohort and IgG antibodies remained present, although at lower levels, for at least 6-8 months after diagnosis.

**Figure 4.**
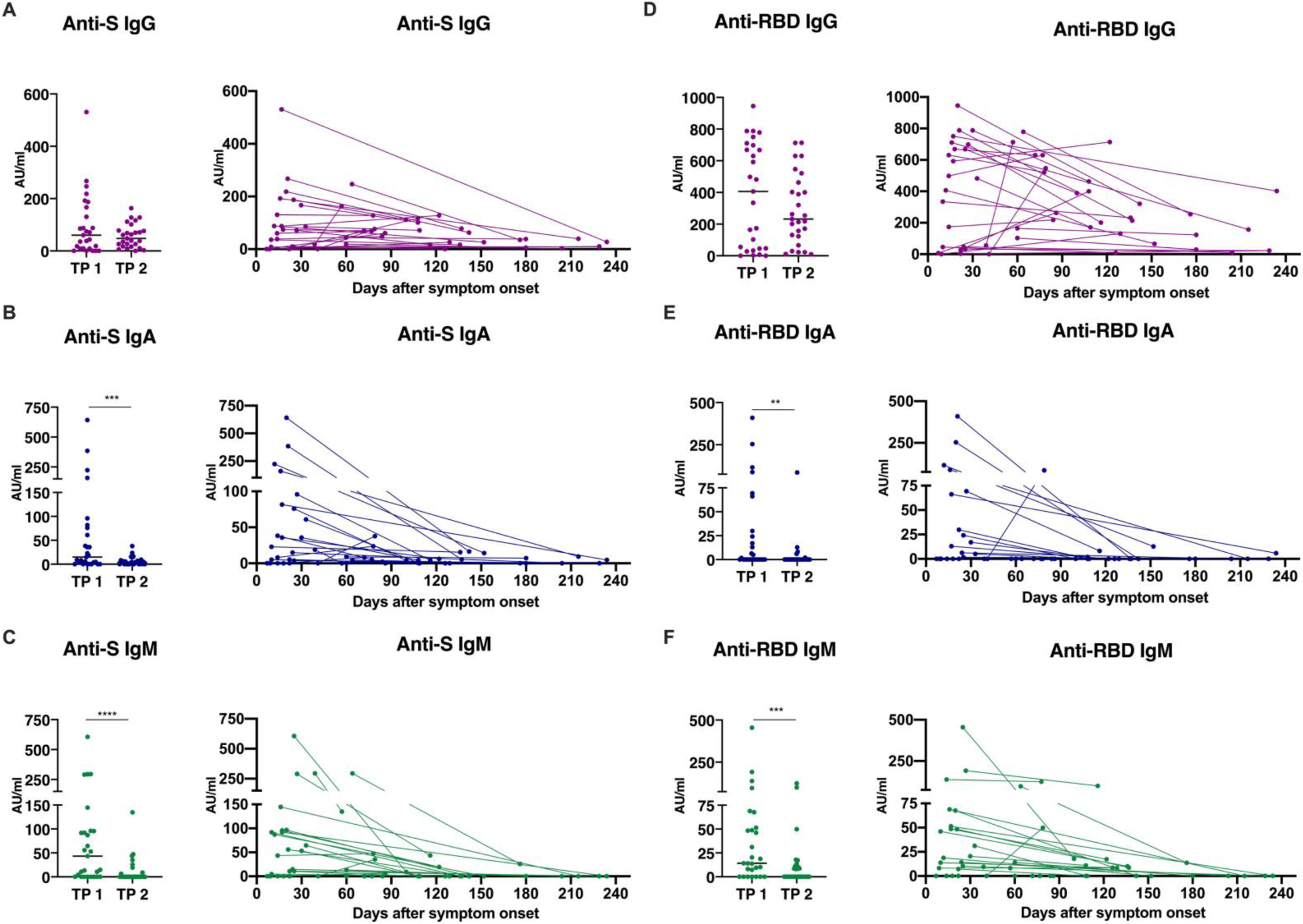
Dynamics of the anti-SARS-CoV-2 antibody levels in paired samples from COVID-19 patients. Levels of anti-S (A, B, C) and anti-RBD (D, E, F) IgG, IgA and IgM antibodies in twenty-seven pairs of COVID-19 patients measured at timepoint 1 (TP1, median day 21, range 7-64) and timepoint 2 (TP2, median day 126, range 57-234) and presented on group (left panel) or individual (right panel) level. Symbols represent individual subjects; horizontal black lines indicate the median. Statistical significance determined by a non-parametric Mann-Whitney U test. **p≤0.001, ***p≤0.001

### Induction of SARS-CoV-2 specific memory B and T cells

To address the question whether SARS-CoV-2-specific memory B and T cells were formed and how long the B- and T-cell mediated responses persist in COVID-19 convalescent individuals, we analyzed 32 PBMC samples collected from 24 patients (mild=11, moderate=4, severe=9, Table S1). No or a negligible number of B cells secreting RBD-specific IgG antibodies were detected in samples from four healthy individuals and seven pre-pandemic buffy coats. Using the highest value observed from all negative controls as a cut-off, RBD-specific IgG producing B cells were detected in 33% (2/6) and 96% (25/26) of the patient samples collected 1-2 month and 3-8 months after onset of symptoms, respectively (Fig. 5A). Thus although the anti-RBD IgG levels declined over time (Fig. 5B), vast majority of the patients developed SARS-CoV-2-specific memory B cells, and these cells remained present in all patients followed up till latest date of study period (n= 13, 6-8 months post symptom onset).

**Figure 5.**
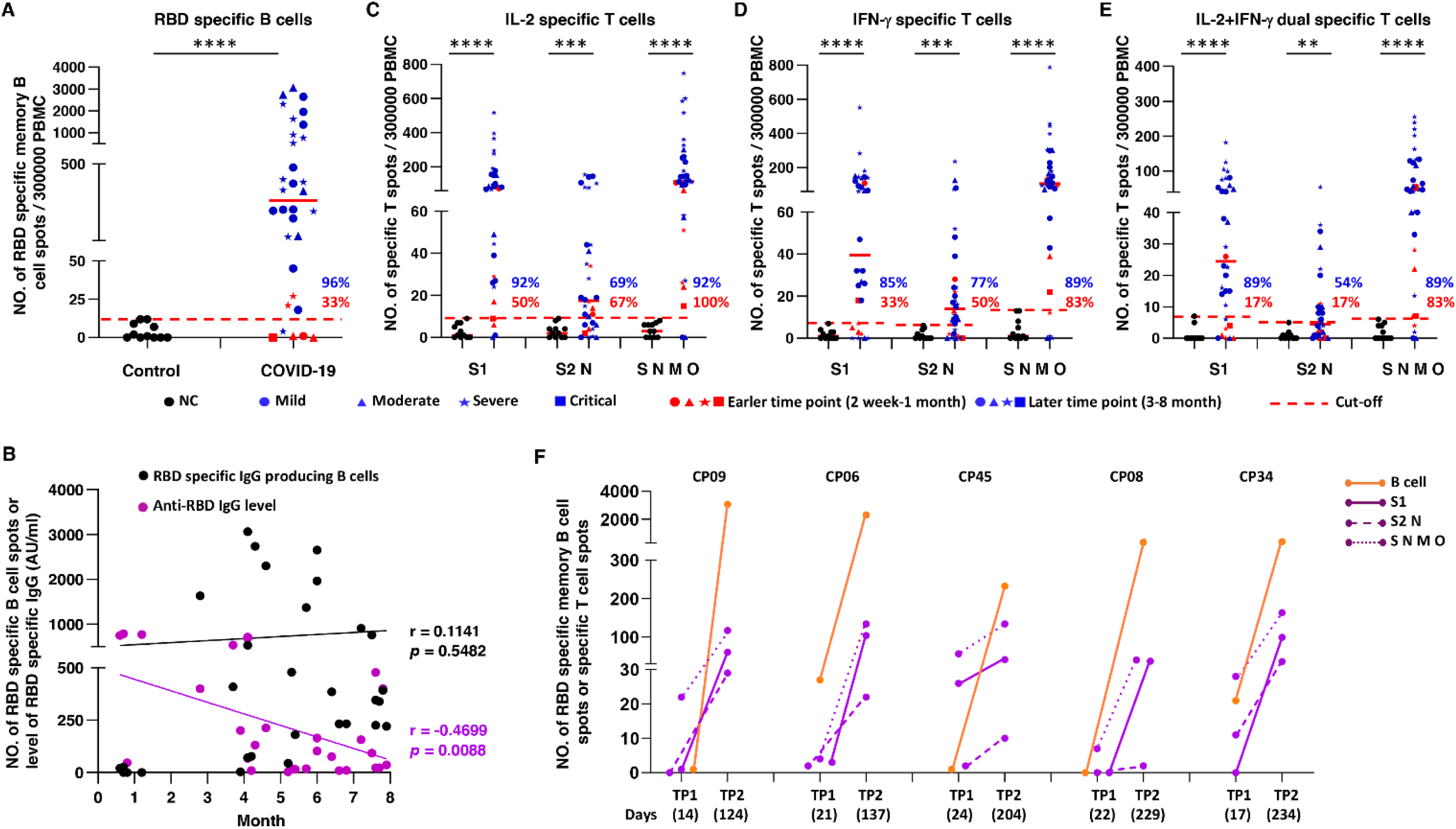
SARS-CoV-2-specific T and memory B cell responses in COVID19 patients. RBD-specific memory B cells from control and COVID-19 patient samples (A). Dynamic of RBD-specific memory B cell and serum anti-RBD IgG levels in COVID-19 patient samples over time (B). SARS-CoV-2-specific T cells specific for the S1, S2N and S N M O protein derived peptides pools and producing IL-2 (**C**), IFN-γ (**D**), or IFN-γ and IL-2 (**E**) in control and COVID-19 patient samples respectively. Increase in specific memory B cells and IFN-γ/IL-2 producing T cells specific for the S1, S2N and S N M O protein derived peptides pools in five patients (with mild (CP45), moderate (CP08, CP09) or severe (CP06, CP34) disease) at time point 1 (TP1) and time point 2 (TP2) (F). The results were expressed as the number of spots per 300 000 seeded cells after subtracting the background spots of the negative control. The red line indicates the median value of the group. The cut-off value was set at the highest number of specific B and T cell spots for the negative controls. P value was calculated by Mann-Whitney U test. **p≤ 0.01, *** p≤0.001, **** p≤0.0001

Furthermore, while no or negligible number of IL-2, IFN-γ or IL-2/IFN-γ producing T cells against three SARS-CoV-2 derived peptide pools were detected in the negative controls, such T cells were observed at a level above the cut-off in 17% to 100% of samples tested 1-2 months after onset of symptoms, depending of the peptide pool tested (S1, S2 N or S N M O protein-derived) and in which cytokines were analyzed (Fig. 5C-E). At later time point (3-8 months after onset of symptoms), T cells specific for the SARS-CoV-2 S1 scanning pool and the defined peptide pool derived from the S, N, M, ORF-3a and ORF-7a proteins were found in 85% to 92% of patient samples, respectively, whereas T cells specific for the S2 N defined peptide pool were observed in 54% to 77% of patient samples (Fig. 5C-E). Overall, T cell response against at least one of the SARS-CoV-2 peptide pools was detectable in all patients (n=6) tested at the early point (1-2 months), and such response maintained in vast majority (96%, 22/23) of patients 3-8 months after onset of symptoms. Notably, the only patient who had no T cell response at 4 months, had a detectable memory B cell response. In two patients, a high number of T cells was also detected in the control test without adding the SARS-CoV-2 peptides, which may suggest an ongoing inflammation.

Moreover, for five paired patient samples collected at an early and later time points (median TP1 = 21 days, median TP2 = 204 days), a significant (p<0.0001) increase in the number of virus-specific B and T cells was observed in a second sample (Fig. 5F). Taken together, SARS-CoV-2-specific memory B and T cells were present in the vast majority of tested convalescent COVID-19 patients, regardless of the initial disease severity, suggesting that the adaptive immunity against SARS-CoV-2 during the natural course of infection is maintained at least for 6-8 months.

## Discussion

In the current study, we measured anti-S and anti-RDB IgG, IgA and IgM antibody levels using normalization against a serially diluted highly positive reference serum pool and by setting a cut-off value based on historical control samples. Our data revealed that an anti-SARS-CoV-2 antibody response was present in a majority of COVID-19 patients as early as two weeks after onset of symptoms, and the level of anti-S and anti-RDB IgG remained stable up to 6 months after diagnosis followed by a decline at month 6-8, while a decrease in anti-S and anti-RDB IgA and IgM levels was observed already between 1 and 3 months after onset of diseases. Our results are in line with previous studies showing a similar longevity and pattern of anti-SARS-CoV-2 antibody response with antibody levels reaching a peak at 23 days following symptom onset and being maintained for at least 4 months ^5 21 22 23 24^, yet contradictory to others, where a low prevalence and rapid decay (within 3 months) of anti–SARS-CoV-2 antibodies in COVID-19 patients with mild or severe disease were observed ^6 25^. In agreement with other reports, we also observed higher anti-RBD IgG antibody titers in men who were more severely affected by SARS-CoV-2 infection while seven percent of patients with a severe disease score in our study did not develop, or had extremely low level of antibodies, after being infected, suggesting that they mounted a weaker anti-viral immune response ^5 7 25^. Higher levels of antibodies in individuals with severe diseases and in male patients might be due to a higher viral load, longer duration of viral shedding ^26 27^ or other host/genetic factors ^28 29^.

Previously reported conflicting findings in prevalence and longevity of anti-SARS-CoV-2 antibody response may result from an absence of a standard assay to measure anti-SARS-CoV-2 antibodies, as a majority of reported studies are based on using different types of SARS-CoV-2 antigens (RBD, S or N proteins). The discrepant results between studies could also be explained by differences between COVID-19 cohorts, as the treatment used, age and sex of study subjects could affect the outcome ^30^. Although the protective role of antibodies against SARS-CoV-2 remains unknown, levels of anti–RBD antibodies seemingly correspond to plasma neutralization activity ^31 24^. While our study cohort is relatively small, the inclusion of Italian and Swedish patients with a different spectrum of disease and the longest follow-up time reported so far (up to 6-8 months), can help to solve the current debate about the persistence of SARS-CoV-2 specific antibodies. It has been reported that antibodies against the two other coronavirus, SARS-CoV and MERS-CoV, could still be detected one to three years after infection onset ^26^, suggesting that SARS-CoV-2 specific antibodies may be present for even longer period than we have observed thus far.

Studies reported until now have mainly been focused on the longevity of the specific antiviral antibody responses. However, the development of memory B and T cell is critical for long-term protection and the longitudinal dynamics of these memory cells remains poorly resolved. Our results show that the majority of patients, irrespective of disease severity, can mount specific memory B- and T-cell responses, which remain present at least 6-8 months post symptom onset. These findings are consistent with preprints showing, using flow cytometry, that RBD-specific memory B cells are generated and maintained up to 3-5 months post-SARS-CoV-2 infection in predominantly mild-moderate cohorts ^32 33 34^. Importantly, we show that these memory B cells are maintained and can secrete RBD-specific IgG antibodies following stimulation.

Previously, it has been shown that S1 and other SARS-CoV-2 proteins derived peptides induce specific T-cell responses in patients with mild to severe disease 1-3 months post symptoms ^1 3 32 35 36^. More recently, it was also reported that SARS-CoV-2-specific T cells are maintained at least 6 months following primary infection in all tested COVID-19 patients with mild to moderated disease ^37^. Our results confirm and extend earlier findings, showing that SARS-CoV-2-specific, IL-2 and/or IFN*-* γ producing cells are present in samples collected from patients with mild to severe diseases, up to 6-8 months post infection. Some studies also noted that T cell responses directed against the S and/or M of SARS-CoV-2 are present in 25-50% of unexposed healthy blood donors, consistent with a high degree of potentially cross-reactive T cell immunity in the general population ^1 36 38^. In our study, the significant increase in number of T cells reacting with SARS-CoV-2-derived peptides from consecutive samples collected at 2-3 weeks and 6-8 months from the same patients strongly suggests a specific response to the SARS-CoV-2 infection, although induction of cross-reactive memory T cells resulting from priming by common cold coronaviruses (e.g. OC43, 229E, NL63 and HKU1) cannot be totally excluded ^39^. The reason we could not detect significant number of cross reactive T cells against the virus might be due to small number of samples tested and/or longer stimulation with the peptides in some other studies ^39^, however cross reactive T cell were also observed in studies using stimulation for a shorter time (9 to 24 hours) ^1 36 38^.

The detection of memory S1-specific T cells marked by production of IL-2, IFN*-* γ and dual production of those cytokine is indicative of induction of T cells with both effector and proliferative potential *in vivo*. IFN*-* γ producing T cells is a hallmark of immunity against intracellular pathogens and although it was not tested in our study, SARS-CoV-2-specific IFN*-* γ producing T cells were previously shown to be of CD4^+^ (Th1-like) or CD8^+^ cytotoxic phenotype ^1 36^. It was shown that convalescent patient donors with undetectable antibodies against the S1 protein of SARS-CoV-2 had T-cell responses more strongly directed against the M than the S1 protein ^37^. Furthermore, a Th1-biased cellular immune response of S-specific IFN*-* γ positive CD4^+^ T cells to pooled S peptides was detected in the majority of monkeys vaccinated with S protein and was associated with induction of specific and neutralizing anti-S antibodies ^40^. Our results suggest that the use of S protein as an immunogen for vaccination has the potential to induce memory T and B cell specific for the S protein and RBD in humans.

Importantly, the decrease in serum IgG antibody levels observed over time in our study was followed by a significant increase in the number of specific memory B and T cells, that could potentially contribute to protection from SARS-CoV-2 reinfection ^32 34^. However, the detection of antibodies to SARS-CoV-2, including neutralizing antibodies, as well as memory B cell and T cell over a long period does not necessarily indicate protective and long-term immunity, and a correlate of protection still needs to be established. Studies on common human coronaviruses show that neutralizing antibodies are induced and reinfections with all seasonal coronaviruses occur usually within three years ^26^. A recent study showed that the duration of protective immunity against common cold coronavirus may last 6 to 12 months ^41^ although others reported that repeat infections are generally associated with milder symptoms and a lower viral load ^26 42^. Single intravenous administration of neutralizing antibodies against the spike protein in outpatients with mild or moderate COVID-19 has been shown to reduce the viral load and reduced length of hospitalization ^43^. Furthermore, infection with SARS-CoV-2 and vaccine against the spike protein can protect rhesus macaque from a challenge infection ^4^. It is thus likely that antibodies and cell-mediated immunity will decrease the risk of reinfection and attenuate the severity in case of reinfection. Characterization of immune responses prior to a known exposure or period of risk is required to identify a correlate of protection ^26^. We thus plan to expand our cohort and follow it over longer intervals of time in order to evaluate the maintenance of immunological memory.

The presence of high level of SARS-CoV-2 specific memory B and T cells in the majority of patients, 6-8 months after infection, suggests that immunity after infection could be at least transiently protective and that development of long-term protective immunity through vaccination might be possible. The discovery of T cell reactivity against S protein epitopes and antibodies against the RBD domain suggests that vaccine development using the S protein to induce antibodies that target RBD is a plausible approach ^35^. To meet the urgent need for SARS-CoV-2 vaccine development, we propose that in addition to analysis of specific antibody responses and their longevity, the development of memory B and T cells, the main components of long-term immunity, as well as correlation with protection from reinfection should be considered.

## Contributors

NS performed the ELISA experiment; A.P., I.C., E.P., A.S., F.Bergami, R.M., H.A., L.D., and F.Ba. contributed to patient data curation, sample preparation and neutralization assay; L.D., M.K.-B. and S.B.-A. designed and/or performed the ELISpot and FluoroSpot experiments; M.V., M.Sa., V.Z., R.B., T.O., and F.M. contributed to the enrollment of patients, patient management and collection of clinical data. J.A., F.Bertoglio, M.Sh. and M.H. performed the production and purification of proteins. N.S., L.D., H.W., M.K.B., M.B.S., H.J., H.A., Y..X, L.H., H.M., and Q.P.-H. contributed to the analysis and interpretation of data. N.S., H.M. and Q.P.-H. drafted the manuscript. H.M., F.B. and Q.P.-H. conceived and supervised the study.

## Acknowledgments

We thank all patients, blood donors and clinicians for their contribution. This project was supported by the European Union’s Horizon 2020 research and innovation programme (ATAC, No 101003650), the Italian Ministry of Health (Ricerca Finalizzata grant no. GR-2013-02358399), CIMED, the Swedish Research Council and funds from the Karolinska Institutet. H.W. was supported by a scholarship from the China Scholarship Council.

