## Supplementary material for "Persistence of SARS-CoV-2 specific B- and T-cell responses in convalescent COVID-19 patients 6-8 months after the infection"

**Table of contents**


**Figure S1** Anti-SARS-CoV-2 antibody responses in COVID-19 patients with different disease scores

**Table S1** Demographic and clinical characteristics of the COVID-19 patients

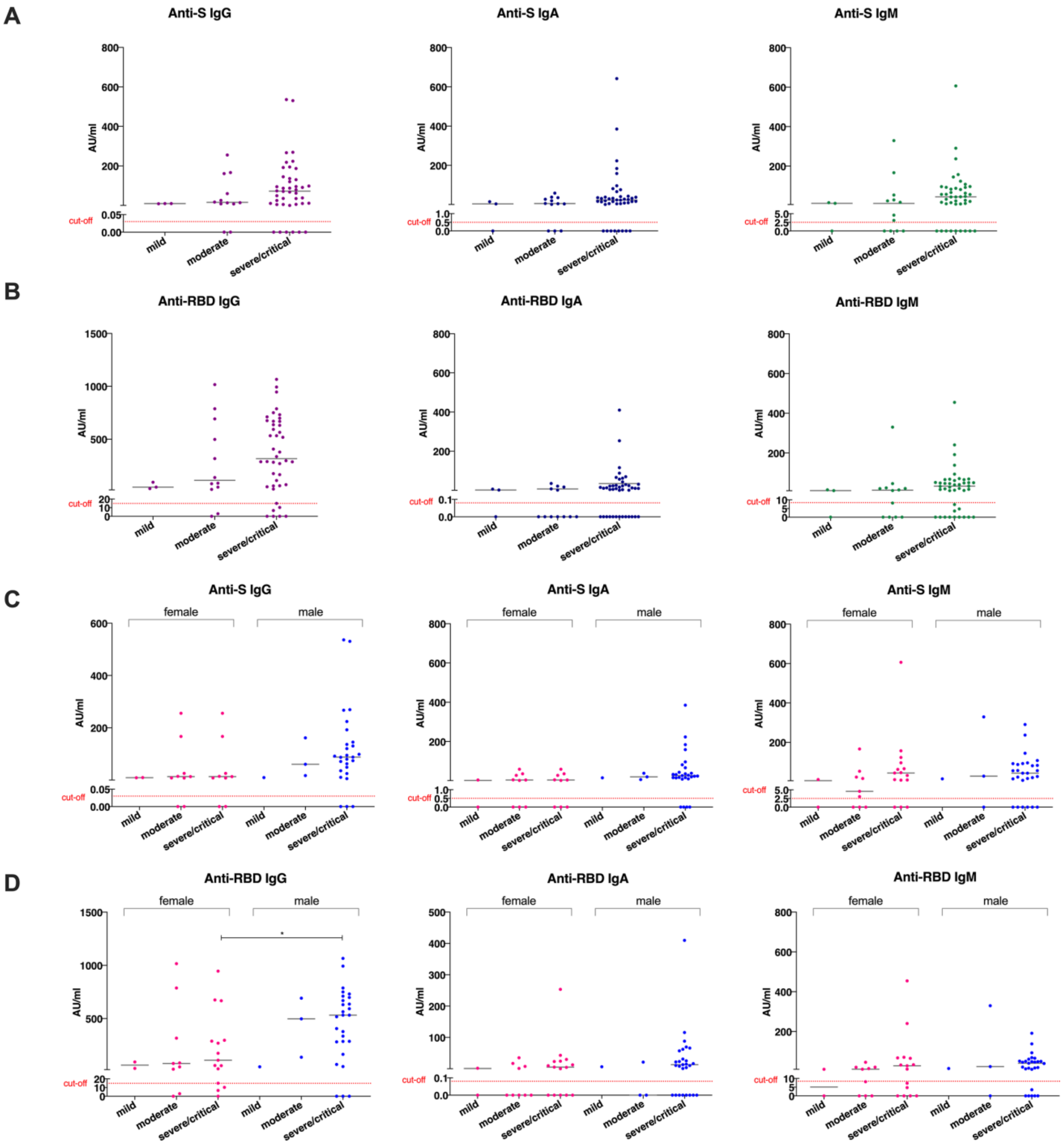


**Figure S1 Anti-SARS-CoV-2 antibody response in COVID-19 patients with different disease scores.** Levels of anti-S and anti-RBD IgG, IgA and IgM antibodies in COVID-19 patients with mild, moderate and severe/critical disease scores (A, B). Levels of anti-S and anti-RBD IgG, IgA and IgM antibodies in female and male COVID-19 patients with mild, moderate and severe/critical disease scores (C, D). Symbols represent individual subjects; horizontal black lines indicate the median. The dashed red line indicates the cut-off value for elevated anti-S and anti-RBD antibody levels (0.03 and 14.81 AU/ml for IgG, 0.5 and 0.08 AU/ml for IgA, 2.5 and 8.4 AU/ml for IgM, respectively) defined using receiver operating characteristic (ROC) curves, based on the antibody responses in historical controls (n=108) and COVID-19 patients (n=55). Statistical significance determined by a non-parametric Mann-Whitney U test. *p=0.0306

### Table S1 Demographic and clinical characteristics

| New ID | Sex | Age | Time point | days after onset of symptoms | ELISA | B ElISpot | T FluroSpot | NT Abs | DiaSorin CLIA results | symptoms | outcome | tumour | heart diseases | hypertension | diabetes | lung diseases | hcv | obesity | immunodeficiency | other comorbidities | other disease or comorbidities (if other comorbidities yes) | Oxygen therapy | ICU | chest CT scan | score (mild, moderate, severe, critical) |
|---|---|---|---|---|---|---|---|---|---|---|---|---|---|---|---|---|---|---|---|---|---|---|---|---|---|
| **Italian cohort** | | | | | | | | | | | | | | | | | | | | | | | | | |
| **CP01** | M | 72 | TP1 | 13 | yes | | | yes | | fever, dyspnea | alive | no | yes | yes | no | no | no | yes | no | no | | VM 40% | no | interstitial pneumonia (low, bilateral) | severe |
| **CP02** | F | 78 | TP1 | 18 | yes | | | yes | | fever, cough | alive | no | yes | yes | no | yes | no | no | no | no | | VM 60% | no | interstitial tickness lobe of the right lung | severe |
| **CP03a** | M | 68 | TP1 | 27 | yes | | | yes | | fever, cough | alive | no | no | no | no | no | no | no | no | no | | CPAP | no | interstitial lung disease (low, bilateral) | severe |
| **CP03b** | | | TP2 | 116 | yes | yes | yes | yes | yes | | | | | | | | | | | | | | | | |
| **CP03c** | | | TP3 | 236 | yes | yes | yes | yes | | | | | | | | | | | | | | | | | |
| **CP04** | M | 73 | TP1 | 23 | yes | | | yes | | fever, cough, dyspnea | death | no | no | no | yes | no | yes | no | no | yes | emophilia A | HFNC | no | na | severe |
| **CP06a** | M | 65 | TP1 | 21 | yes | yes | yes | yes | | fever, cough, dyspena, asthenia | alive | no | no | no | yes | no | no | no | no | no | | VM 50% | no | bronchovascular thickening and opacity | severe |
| **CP06b** | | | TP2 | 137 | yes | yes | yes | yes | yes | | | | | | | | | | | | | | | | |
| **CP06c** | | | TP3 | 226 | yes | yes | yes | yes | yes | | | | | | | | | | | | | | | | |
| **CP07a** | F | 61 | TP1 | 25 | yes | | | yes | | fever, cough | alive | na | na | na | na | na | na | na | na | na | | no | no | na | mild |
| **CP07b** | | | TP2 | 229 | yes | yes | yes | yes | yes | | | | | | | | | | | | | | | | |
| **CP08a** | F | 72 | TP1 | 22 | yes | yes | yes | | | fever, cough, diarrhea | alive | no | no | yes | no | no | no | no | no | no | | no | no | lung opacity basal bilateral | moderate |
| **CP08b** | | | TP2 | 229 | yes | yes | yes | yes | yes | | | | | | | | | | | | | | | | |
| **CP09a** | M | 56 | TP1 | 14 | | yes | yes | yes | | fever | alive | no | no | yes | no | no | no | no | no | no | | no | no | bronchovascular thickening | moderate |
| **CP09b** | | | TP2 | 124 | yes | yes | yes | yes | | | | | | | | | | | | | | | | | |
| **CP10a** | F | 32 | TP1 | 10 | yes | | | | | fever, cough, myalgia | alive | no | no | no | no | no | no | no | no | no | | no | no | no lung densification | moderate |
| **CP10b** | | | TP2 | 126 | yes | yes | yes | yes | yes | | | | | | | | | | | | | | | | |
| **CP11a** | F | 44 | TP1 | 9 | yes | | | | | fever, diarrhea | alive | no | no | no | no | no | yes | no | no | no | | no | no | no lung densification | moderate |
| **CP11b** | | | TP2 | 129 | yes | yes | yes | | yes | | | | | | | | | | | | | | | | |
| **CP12** | M | 70 | TP1 | 23 | yes | | | | | fever, cough, dyspena | alive | no | no | no | yes | no | no | yes | no | no | | HFNC | no | bronchovascular thickening | severe |
| **CP13a** | F | 70 | TP1 | 30 | yes | | | yes | | cough | alive | no | yes | yes | no | no | no | no | no | no | | no | no | na | moderate |
| **CP13b** | | | TP2 | 108 | yes | | | yes | | | | | | | | | | | | | | | | | |
| **CP14** | M | 78 | TP1 | 12 | yes | | | | | fever,asthenia | death | yes | no | yes | yes | no | no | no | no | yes | previous colon carcinoma | VM 60% | no | interstitial lung disease | severe |
| **CP15** | F | 86 | TP1 | 18 | yes | | | yes | | fever, cough | alive | no | yes | yes | yes | no | no | no | no | no | | no | no | na | moderate |
| **CP16a** | F | 50 | TP1 | 25 | yes | | | | | fever, dyspnea, | alive | no | no | yes | no | no | no | no | no | no | | VM 60% | no | lung densification | severe |
| **CP16b** | | | TP2 | 100 | yes | | | yes | yes | | | | | | | | | | | | | | | | |
| **CP17a** | M | 42 | TP1 | 10 | yes | | | yes | | fever, cough, dyspnea | alive | no | no | no | no | no | no | no | no | no | | LFNC | no | bronchovascular thickening | severe |

| | | | | | | | | | | | | | | | | | | | | | | | | | |
|---|---|---|---|---|---|---|---|---|---|---|---|---|---|---|---|---|---|---|---|---|---|---|---|---|---|
| **CP17b** | | | TP2 | 136 | yes | | | yes | yes | | | | | | | | | | | | | | | | |
| **CP18** | M | 53 | TP1 | 18 | yes | | | | | fever, dispnea | alive | no | no | no | no | no | no | no | no | no | | VM 60% | no | bilateral densification (medium-low area) | severe |
| **CP19a** | M | 77 | TP1 | 17 | yes | | | | | fever, cough | alive | no | yes | no | no | no | no | no | no | yes | gatroesophageal reflux, renal disfunction | HFNC | no | bilateral densification | severe |
| **CP19b** | | | TP2 | 122 | yes | yes | yes | yes | yes | | | | | | | | | | | | | | | | |
| **CP19c** | | | TP3 | 228 | yes | yes | yes | yes | yes | | | | | | | | | | | | | | | | |
| **CP20** | M | 57 | TP1 | 20 | yes | | | | | fever, cough, dyspnea | alive | no | no | no | no | no | no | no | no | no | | VM 40% | no | lung densification (low, bilateral) | severe |
| **CP21** | M | 81 | TP1 | 28 | yes | | | yes | | fever, cough | alive | no | no | yes | no | no | no | no | no | no | | VM 35% | no | multiple bilateral densification | severe |
| **CP23** | M | 49 | TP1 | 15 | yes | | | yes | | fever, dyspnea | alive | no | no | no | no | no | no | no | no | no | | HFNC | no | bilateral lung densification | severe |
| **CP24** | F | 60 | TP1 | 39 | yes | | | yes | | fever, dyspnea | alive | no | no | yes | no | no | no | no | no | no | | HFNC | no | multiple bilateral densification | severe |
| **CP25** | F | 62 | TP1 | 35 | yes | | | yes | | fever, diarrhea | alive | no | no | yes | no | no | no | no | no | yes | osteoporosis, anemia, previous anticoagulant lupus | no | no | interstitial lung densification (peripheral basal area, right) | moderate |
| **CP26** | F | 55 | TP1 | 35 | yes | yes | yes | yes | | fever, cough | alive | no | no | no | no | no | no | yes | no | no | | HFNC/CPAP | yes | bronchovascular thickening | critical |
| **CP27** | F | 72 | TP1 | 26 | yes | | | | | fever, cough, dyspnea | alive | yes | no | yes | no | no | yes | no | no | yes | anostrazole therapy for previous mammalian carcinoma | HFNC/CPAP | no | bilateral interstitial opacity | severe |
| **CP28** | F | 50 | TP1 | 30 | yes | | | yes | | fever, cough | alive | no | no | yes | no | no | no | yes | no | yes | microcytic anemia | HFNC/CPAP | no | irregular densification | severe |
| **CP29** | F | 53 | TP1 | 23 | yes | | | yes | | fever, dyspnea | alive | no | no | yes | no | no | yes | no | no | no | | HFNC/CPAP | yes | bronchovascular thickening | critical |
| **CP30** | M | 72 | TP1 | 27 | yes | | | yes | | fever, dyspnea | alive | no | no | yes | yes | no | no | no | no | yes | hypothyroidism, previous prostatic carcinoma | HFNC | no | bilateral densification (basal area) | severe |
| **CP33** | M | 45 | TP1 | 19 | yes | | | yes | | fever,cough | alive | no | no | yes | no | yes | no | no | no | no | | CPAP | no | lung opacity | severe |
| **CP34a** | M | 67 | TP1 | 17 | yes | yes | yes | | yes | fever | alive | no | yes | yes | no | no | no | no | no | yes | previous pneumococcical pneumonia | HFNC | no | parenchimal densification (low basal area left) | severe |
| **CP34b** | | | TP2 | 234 | yes | yes | yes | yes | yes | | | | | | | | | | | | | | | | |
| **CP35a** | M | 71 | TP1 | 12 | yes | yes | yes | | | fever, cough, dyspnea | alive | no | no | yes | no | no | no | no | no | no | | HFNC | no | bilateral interstitial opacity | severe |
| **CP35b** | | | TP2 | 152 | yes | yes | | yes | yes | | | | | | | | | | | | | | | | |
| **CP36** | M | 84 | TP1 | 8 | yes | | | yes | | fever, cough, dyspnea | alive | no | no | yes | no | no | no | no | no | no | | no | no | na | moderate |
| **CP37a** | F | 43 | TP1 | 20 | yes | | | | | fever, cough, dyspnea | alive | no | no | yes | yes | no | no | yes | no | no | | HFNC | no | bronchovascular thickening | severe |
| **CP37b** | | | TP2 | 142 | yes | | | | yes | | | | | | | | | | | | | | | | |
| **CP38** | F | 89 | TP1 | 11 | yes | | | yes | | fever, cough, dyspnea | death | yes | yes | yes | no | no | no | no | no | no | | na | no | na | severe |
| **CP39** | M | 37 | TP1 | 35 | yes | | | yes | | fever, cough, dyspena | alive | no | no | no | no | no | no | no | no | no | | HFNC/CPAP | yes | bronchovascular thickening | critical |
| **CP40** | F | 67 | TP1 | 19 | yes | | | | | fever, cough | alive | no | no | no | no | no | no | no | no | yes | hypothyroidism | CPAP | no | interstitial lung disease and parenchimal densification (left), | severe |

| | | | | | | | | | | | | | | | | | | | | | | | | | |
|---|---|---|---|---|---|---|---|---|---|---|---|---|---|---|---|---|---|---|---|---|---|---|---|---|---|
| **CP41** | M | 57 | TP1 | 10 | yes | | | yes | | fever, cough, diarrhea | alive | no | no | yes | no | no | no | no | no | yes | not specified | CPAP | no | na | severe |
| **CP42** | M | 77 | TP1 | 19 | yes | | | yes | | fever, dyspnea | alive | no | yes | yes | yes | no | no | no | no | no | | HFNC/CPAP | no | interstitial lung disease (medium-low, bilateral) | severe |
| **CP43** | M | 69 | TP1 | 13 | yes | | | yes | | fever, cough, dyspena | alive | no | no | yes | yes | no | no | no | no | no | | CPAP | yes | interstitial lung disease | critical |
| **CP44a** | M | 60 | TP1 | 16 | yes | | | yes | | fever, dyspena | alive | no | no | no | no | no | no | no | no | yes | peripheral artherial disease | HFNC | no | bilateral lung opacity | severe |
| **CP44b** | | | TP2 | 215 | yes | yes | yes | yes | | | | | | | | | | | | | | | | | |
| **CP45a** | M | 53 | TP1 | 24 | yes | yes | yes | yes | | fever | alive | na | na | na | na | na | na | na | na | na | | no | no | na | mild |
| **CP45b** | | | TP2 | 204 | yes | yes | yes | yes | yes | | | | | | | | | | | | | | | | |
| **CP46a** | F | 80 | TP1 | 26 | | | | | yes | fever, diarrhea | alive | no | no | yes | no | no | yes | no | no | yes | gastroesophageal reflux disease/osteoporosis | HFNC | no | interstitial lung disease | severe |
| **CP46b** | | | TP2 | 67 | yes | | | | | | | | | | | | | | | | | | | | |
| **CP47a** | F | 59 | TP1 | 26 | yes | | | yes | | fever, dyspnea | alive | no | no | no | no | no | no | yes | no | yes | depressive syndrome | HFNC | no | lung opacity | severe |
| **CP47b** | | | TP2 | 35 | | | | | | | | | | | | | | | | | | | | | severe |
| **CP48a** | F | 60 | TP1 | 39 | yes | | | yes | | fever cough, dyspnea | alive | no | yes | yes | no | no | no | no | no | yes | not specified | CPAP/PEEP | no | lung opacity (medium left area) | severe |
| **CP48b** | | | TP2 | 57 | yes | | | | | | | | | | | | | | | | | | | | |
| **CP49a** | M | 50 | TP1 | 21 | yes | | | yes | | fever, cough | alive | no | no | no | no | no | no | no | no | no | | VM 40% | no | bilateral densification | severe |
| **CP49b** | | | TP2 | 42 | | | | | | | | | | | | | | | | | | | | | severe |
| **CP50a** | M | 71 | TP1 | 21 | yes | | | yes | | fever, dyspnea | alive | no | no | yes | no | no | no | no | no | yes | prostatic hyperplasia | HFNC | no | lung bilateral densification | severe |
| **CP50b** | | | TP2 | 42 | | | | | | | | | | | | | | | | | | | | | severe |
| **CP51** | M | 55 | TP1 | 26 | yes | | | yes | | fever, dyspnea | alive | no | no | no | no | no | no | no | no | no | | HFNC, CPAP | yes | multiple lung densification | critical |
| **CP52a** | M | 73 | TP1 | 41 | yes | | | | yes | fever, cough | alive | no | no | no | no | no | yes | no | no | yes | prostatic hyperplasia, gladbladder stones | HFNC | no | parenchymal densification (right lung, low area) | severe |
| **CP52b** | | | TP2 | 79 | yes | | | | | | | | | | | | | | | | | | | | |
| **CP53a** | M | 61 | TP1 | 18 | yes | | | | yes | fever, cough, dyspnea | alive | no | no | yes | no | no | no | no | no | no | | CPAP | no | lung densification | severe |
| **CP53b** | | | TP2 | 72 | yes | | | | | | | | | | | | | | | | | | | | |
| **CP54a** | M | 66 | TP1 | 14 | yes | | | | yes | fever, cough, dyspena | alive | no | no | no | no | no | no | no | no | yes | not specified | no | no | na | moderate |
| **CP54b** | | | TP2 | 77 | yes | | | | | | | | | | | | | | | | | | | | |
| **CP55a** | F | 76 | TP1 | 14 | yes | | | yes | | fever, cough | alive | no | no | yes | no | no | yes | no | no | no | | LFNC | no | lung opacity (medium-low left region) | severe |
| **CP55b** | | | TP2 | 84 | yes | | | | | | | | | | | | | | | | | | | | |
| **CP56a** | M | 69 | TP1 | 14 | yes | | | | yes | cough, dyspena, asthenia | alive | no | no | no | no | yes | no | no | no | yes | psoriatic arthritis | LFNC | no | bronchovascular thickening (medium-low right) | severe |
| **CP56b** | | | TP2 | 78 | yes | | | | | | | | | | | | | | | | | | | | |
| **CP57a** | F | 72 | TP1 | 22 | yes | | | | yes | fever, cough, dyspnea | alive | no | no | yes | no | no | no | no | no | yes | dyslipidemia | VM 60% | no | multiple lung foci | severe |
| **CP57b** | | | TP2 | 108 | yes | | | | | | | | | | | | | | | | | | | | |
| **CP58a** | M | 74 | TP1 | 33 | yes | | | | yes | fever, cough | alive | yes | no | yes | no | no | no | no | no | yes | Paget syndrome | CPAP | no | densification medium-high left lung, opacity medium-high right lung | severe |
| **CP58b** | | | TP2 | 109 | yes | | | | | | | | | | | | | | | | | | | | |

| | | | | | | | | | | | | | | | | | | | | | | | | | |
|---|---|---|---|---|---|---|---|---|---|---|---|---|---|---|---|---|---|---|---|---|---|---|---|---|---|
| **CP59** | F | 37 | TP1 | 18 | yes | | | yes | yes | fever | alive | na | na | na | na | na | na | na | na | na | | no | no | na | mild |
| **CP60a** | F | 54 | TP1 | 7 | yes | | | yes | | fever, dyspnea, diarrhea, anosmia | alive | no | no | no | no | no | no | no | no | no | | VM | no | bilateral parenchymal opacity | severe |
| **CP60b** | | | TP2 | 86 | yes | | | yes | yes | | | | | | | | | | | | | | | | |
| **CP61** | M | 43 | TP1 | 118 | yes | | | yes | yes | fever, dyspnea | alive | no | no | no | no | no | no | no | no | yes | gastroesophageal reflux, kidney stones | VM/HFNC, CPCP | no | lung densification (basal right and left) | severe |
| **CP62** | M | 49 | TP1 | 110 | yes | | | yes | yes | cough | alive | no | no | no | yes | no | no | yes | no | yes | ovarian cystis and uterine fibroid | no | no | lung opacity (bilateral) | moderate |
| **CP63** | M | 32 | TP1 | 85 | yes | yes | yes | yes | yes | dyspnea | alive | no | no | no | no | no | no | no | no | yes | gastritis (pump inhinitors treatment) | HFNC | no | bronchovascular thickening | severe |
| **CP64** | F | 51 | TP1 | 107 | yes | | | yes | yes | fever, cough | alive | no | no | no | no | no | no | no | no | no | | HFNC | no | lung densification | severe |
| **CP65a** | M | 65 | TP1 | 64 | yes | | | yes | yes | fever | alive | no | no | no | no | no | no | no | no | no | | VM/ CPAP | no | bilateral lung opacity and initial lung densification | severe |
| **CP65b** | | | TP2 | 176 | yes | | | yes | yes | | | | | | | | | | | | | | | | |
| **CP66** | M | 56 | TP1 | 93 | yes | | | yes | yes | fever, dyspnea | alive | no | no | no | no | yes | no | yes | no | yes | hypothyroidism | PEEP/ CPAP-VM | no | densification medium-low right and low left lung | severe |
| **CP67** | M | 52 | TP1 | 89 | yes | | | yes | yes | fever, dyspnea, anosmia | alive | no | no | no | yes | no | no | yes | no | no | | HFNC | no | pleural effusion (basal right lung) | severe |
| **CP68a** | M | 64 | TP1 | 101 | yes | | | yes | yes | fever, cough, dyspnea | alive | no | no | no | no | no | no | no | no | no | | no | no | na | moderate |
| **CP68b** | | | TP2 | 210 | yes | | | yes | yes | | | | | | | | | | | | | | | | |
| **CP69** | F | 56 | TP1 | 112 | yes | yes | yes | yes | yes | fever, cough, dyspnea | alive | no | no | yes | no | no | no | no | no | no | | CPAP | no | bronchovascular thickening | severe |
| **CP70** | M | 52 | TP1 | 122 | yes | yes | yes | yes | yes | fever, cough, hypoxia | alive | no | no | yes | no | no | no | no | no | no | | VM | no | medium-upper right and medium left parenchymal densification | severe |
| **CP81** | F | 88 | TP1 | 11 | yes | | | yes | yes | fever | alive | no | no | yes | no | no | no | yes | no | yes | dyslipidemia, renal cystis, osteoporosis, stroke | no | no | no | moderate |
| **CP82** | F | 79 | TP1 | 15 | yes | | | yes | yes | fever, dyspnea | alive | no | no | yes | yes | no | yes | no | no | yes | ostheoarthritis | VM60% | no | bilateral densification low-medium area | severe |
| **CP83** | F | 48 | TP1 | 7 | yes | | | yes | yes | fever | alive | no | no | yes | no | no | no | no | no | yes | hypothiroidism, diverticulitis (sigmoid) | no | no | absence of evident lesions | moderate |
| **CP84** | M | 83 | TP1 | 15 | yes | | | yes | yes | asthenia, diarrhea | alive | no | yes | yes | yes | no | no | no | no | yes | chronic anemia, cronic renal disease | no | no | bronchovascular thickening, dx basal lung opacity | moderate |
| **CP85** | F | 53 | TP1 | 10 | yes | | | yes | yes | fever, asthenia | alive | no | no | no | no | no | no | no | no | no | | no | no | multiple bilateral densification | moderate |
| **CP86** | F | 73 | TP1 | 10 | yes | | | yes | yes | fever | alive | no | no | no | no | no | no | no | no | no | | HFNC | no | initial lung densification | severe |
| **CP87** | M | 34 | TP1 | 196 | yes | | | yes | yes | fever, asthenia, anosmia | alive | no | no | no | no | no | no | no | no | no | | no | no | no | mild |
| **CP88** | F | 81 | TP1 | 26 | yes | | | yes | yes | fever, asthenia | alive | no | yes | no | yes | no | no | no | no | yes | transient ischemic attack | no | no | no lung densification | moderate |
| **CP89** | M | 57 | TP1 | 21 | yes | | | yes | yes | fever, dyspnea | alive | no | no | yes | yes | no | no | no | no | yes | renal disfunction | VM40% | no | multiple bilateral densification | severe |
| **CP90** | F | 73 | TP1 | 22 | yes | | | yes | yes | fever, cough | alive | no | yes | yes | yes | no | no | no | no | yes | Sjogren syndrom, previous B lymphoma | HFNC | no | bilateral densification (basal area) | severe |
| **CP91** | M | 66 | TP1 | 206 | yes | | | yes | yes | fever, asthenia | alive | no | no | no | no | no | no | no | no | no | | no | no | no | mild |

| | | | | | | | | | | | | | | | | | | | | | | | | | |
|---|---|---|---|---|---|---|---|---|---|---|---|---|---|---|---|---|---|---|---|---|---|---|---|---|---|
| **CP92** | M | 80 | TP1 | 9 | yes | | | yes | yes | fever, asthenia, cough | alive | no | no | yes | no | no | no | no | no | yes | renal disease, depression | VM | no | multiple bilateral densification | severe |
| **Swedish cohort** | | | | | | | | | | | | | | | | | | | | | | | | | |
| **CP31a** | F | 75 | TP1 | 60 | yes | | | | | na | alive | no | na | na | na | na | na | na | na | na | na | na | no | na | mild |
| **CP31b** | | | TP2 | 180 | yes | yes | yes | | | | | | | | | | | | | | | | | | |
| **CP71** | M | 29 | TP1 | 170 | yes | yes | yes | | | myalgia, headache, anosmia,asthenia,chest pain | alive | no | na | na | na | na | na | na | na | na | na | na | no | na | mild |
| **CP72a** | M | 65 | TP1 | 60 | yes | | | | | fever | alive | no | na | na | na | na | na | na | na | na | na | na | no | na | mild |
| **CP72b** | | | TP2 | 180 | yes | yes | yes | | | | | | | | | | | | | | | | | | |
| **CP73** | M | 48 | TP1 | 180 | yes | yes | yes | | | fever, cough | alive | no | na | na | na | na | na | na | na | na | na | na | no | na | mild |
| **CP74** | F | 61 | TP1 | 198 | yes | yes | yes | | | cough, myalgia, asthenia, anosmia | alive | no | na | na | na | na | na | na | na | na | na | na | no | na | mild |
| **CP75** | M | 56 | TP1 | 193 | yes | yes | yes | | | cough, myalgia, asthenia, anosmia | alive | no | na | na | na | na | na | na | na | na | na | na | no | na | mild |
| **CP76** | F | 29 | TP1 | 219 | yes | yes | yes | | | cough | alive | no | na | na | na | na | na | na | na | na | na | na | no | na | mild |
| **CP77** | F | 43 | TP1 | 238 | yes | | | | | asthenia | alive | no | na | na | na | na | na | na | na | na | na | na | no | na | mild |
| **CP78** | M | 55 | TP1 | 157 | yes | yes | yes | | | fever, myalgia, asthenia, anosmia | alive | no | na | na | na | na | na | na | na | yes | kidney disease | na | no | na | mild |
| **CP79** | F | 52 | TP1 | 160 | yes | yes | yes | | | fever, myalgia,asthenia, anosmia | alive | no | na | na | na | na | na | na | na | na | na | na | no | na | mild |

| | | | | | | | | |
|---|---|---|---|---|---|---|---|---|
| **NC1** | na | na | | | yes | yes | yes | pre-pandemic buffy-coat control collected in 2020 Jan. |
| **NC2** | na | na | | | | yes | yes | pre-pandemic buffy-coat control collected in 2011. |
| **NC3** | na | na | | | | yes | yes | pre-pandemic buffy-coat control collected in 2017. |
| **NC4** | na | na | | | | yes | yes | pre-pandemic buffy-coat control collected in 2019 Oct. |
| **NC5** | na | na | | | | yes | yes | pre-pandemic buffy-coat control collected in 2019 Oct. |
| **NC6** | na | na | | | | yes | yes | pre-pandemic buffy-coat control collected in 2019 Oct. |
| **NC7** | na | na | | | | yes | yes | pre-pandemic buffy-coat control collected in 2020 Jan. |
| **NC8** | F | 50 | | | yes | yes | yes | |
| **NC9** | M | 39 | | | yes | yes | yes | |
| **NC10** | F | 59 | | | | yes | yes | |
| **NC11** | M | 42 | | | | yes | yes | |

**M** Male
**F** Female
**TP1** Time point 1
**TP2** Time point 2
**na** Not available
**NT Abs** SARS-CoV-2 microneutralization assay (Bonelli et al. 2020 J Clin Microbiol 2020;58(9):e01224-20)
**hcv** Hepatitis C virus
**VM** Venturi oxygen mask
**CPAP** Continuous positive airway pressure
**HFNC** High-flow nasal cannula
**CLIA** Chemiluminescence immunoassay (Bonelli et al. 2020 J Clin Microbiol 2020;58(9):e01224-20)
**ICU** Intensive care unit
**CT** Computed tomography
